# Glucosidase inhibitors suppress SARS-CoV-2 in tissue culture and may potentiate

**DOI:** 10.1101/2021.05.14.444190

**Authors:** Hanako Reyes, Yanming Du, Tianlun Zhou, Xuping Xie, Pei Yong Shi, Susan Weiss, Timothy M. Block

## Abstract

Iminosugar glucosidase inhibitors prevent the folding of a range of viral N-linked glycoproteins, ranging from hepatitis B to Ebola. We recently showed they inhibit folding and function of the ACE2 protein, which is the receptor for SARS-CoV-2, and they have also inhibited the SARS Spike polypeptides. Here we report that the imino sugar glucosidase inhibitors, N-butyl deoxynojirimycin (NBDNJ), which is approved for management of lysosomal storage disease (sold as Zavesca), and ureido-N-hexyl deoxynojirimycine (BSBI-19029), suppress the replication of SARS-ncCoV-2/USA/WA1/2020 strain, in tissue culture. Moreover, combinations of either of these iminosugars with Remdesivir were particularly potent in suppressing SARS-CoV-2. Briefly, NBDNJ, 19029 and Remdesivir suppressed SARS-CoV-2 production in A549^ACE2^ human lung cells with IC90s of ~130 μM, ~4.0 μM, and 0.006 μM respectively. The combination of as little as 0.037 μM of NBDNJ or 0.04 μM 19029, respectively and 0.002 μM Remdesivir yielded IC90s. Medical strategies to manage SARS-CoV-2 infection of people are urgently needed, and although Remdesivir and Favipiravir have shown efficacy, it is limited. NBDNJ was recently reported by others to have tissue culture activity against SARS-CoV-2, so our report confirms this, and extends the findings to a more potent iminosugar, 19029 and combination with Remdesivir. Since both NBDNJ and Remdesivir are both approved and available for human use, the possibility that NBDNJ has mono therapeutic value against SARS-CoV-2 as well as can enhance Remdesivir, may have clinical implications, which are discussed, here.

## INTRODUCTION

Severe acute respiratory syndrome coronavirus 2 (SARS-CoV-2) is the etiological agent of the Coronavirus disease of 2019 (COVID-19). The WHO has declared it a public health emergency of international concern (Cucinotta et al., 2020; Ghebreyesus, 2020). Since its emergence in December 2019, and its isolation on January 7, 2020, the number of infections has grown explosively (Hoehl et al., 2020; Holshue et al., 2020; Lipsitch., et al 2020). As of April 10, 2021, there are > 40 million reports of infection with more than 550,000 deaths in the US alone, up from 600 cases, with 21 deaths March 1, 2020 (Engineering JHCfSSA, CDC Covid Data tracker, Angulo et al., 2021).

Antivirals effective against SARS-CoV-2 and medical management of COVID-19 are urgently needed. There are currently, not surprisingly given its recent discovery, no therapeutic drugs approved for use against SARS-CoV-2 outside of emergency authorizations. Like Middle East respiratory syndrome coronavirus (MERS-CoV) and severe acute respiratory syndrome (SARS-CoV), and 229E, NL63, OC43, HKU1, SARS-CoV-2 is a member of the Coronavirus family (Weiss & Navas-Martin, 2005). All strains have ~30kb of positive stranded RNA genomes (Weiss & Navas-Martin, 2005; Walls et al., 2020; Graham et al., 2008; Andersen, et al., 2020). Among the ~25-30 polypeptides encoded by its RNA genome are a nucleocapsid protein (N), a Spike (S), Membrane (M) and small membrane glycoprotein (E) and virally encoded RNA dependent RNA polymerases (RdRp), which mediates replication of genome RNA and transcription of mRNAs. The transmembrane spike glycoprotein (S) mediates binding to a receptor on host cells, in the case of SARs-CoV-2, angiotensin converting enzyme 2 (ACE2) (Cuervo & Grandvaux, 2020; Li et al., 2003; Lu et al., 2020; Walls et al., 2021; Wan et al., 2020).

The Gilead prodrug nucleoside (Nuc), GS5734 (Remdesivir), was shown to have activity against SARS-CoV in vitro, in the low to sub micromolar range (Tchesnokov et al., 2020; Simonis et al., 2021). Its target is the viral nsp12 RdRp (Agostini et al., 2018) and in animal models and human clinical trials, Remdesivir improved outcome (Beigel et al., 2020; Spinner et al., 2021). Remdesivir now has US FDA Emergency Use Authorization for COVID-19

We have been working with a family of iminosugars, we call iminovirs (examples in Fig. 1). These have Deoxynorimycin (DNJ) head groups and have been shown to have antiviral activity against a number of “enveloped” viruses (Mehta et al., 1998). The molecular basis for how the iminosugars inhibit binding and morphogenesis of these viruses is well established. They are competitive inhibitors of the host Endoplasmic Reticular (ER) glucosidases I & II which are critical for the Calnexin dependent folding of specific N-glycosylated proteins (Mehta et al., 1998; Schrag et al., 2001). We (and others) found, 2 decades ago, that most cell glycoproteins can use alternatives to the ER glucosidases for their folding and secretion(Block et al., 1994; Dwek et al., 2002; Chang et al., 2013), but a subset of viral and cellular glycoproteins appear to be very dependent upon them (Chang et al., 2013, 2015; Ma et al., 2018).

**Figure 1:**
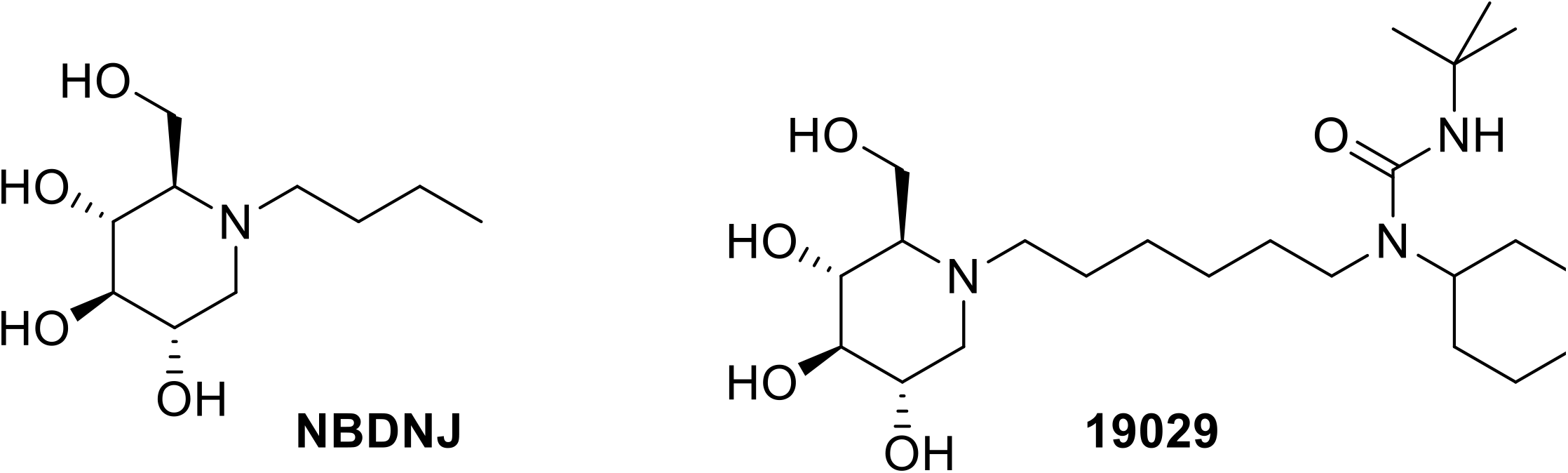
Structure of iminosugar compounds NBDNJ and BSBI-19029.

More recently we reported that maturation of the ACE2 receptor, unlike most cellular glycoproteins, is very sensitive to ER glucosidase mediated folding (Zhao et al., 2015). Moreover, we also showed that cell entry of pseudostyle virus whose entry is dependent upon the SARS-CoV spike polypeptide, mediated by the SARS-CoV protein, was greatly reduced in the presence of glucosidase inhibitors, and this was a function of inhibition of ACE 2 (Zhao et al., 2015). Others also reported (Fukushi et al., 2012) SARS-CoV was inhibited by our glucosidase inhibitors.

Since SARS-CoV-2 and SARS-CoV both use the ACE2 receptor, and both specify essential highly N-glycosylated spike polypeptides, we reasoned SARS-CoV-2 would also be sensitive to ER glucosidase inhibitions. Also, since mechanisms of antiviral inhibition of the glucosidase inhibitors and Remdesivir and Favipiravir are distinct, we reasoned the glucosidase inhibitors would be able to be additive or potentiate the anti-SARS-CoV-2 activity of direct antiviral acting drugs such as Remdesivir.

Here we report that the iminosugar glucosidase inhibitors NBDNJ (“Miglustat/Zavesca”) and ureido-N-hexyl deoxynojirimycin (BSBI-19029) have significant ability to repress a SARS-CoV-2 virus, in tissue culture, at concentrations at least 100 times less than their cytotoxic concentrations. Moreover, as expected, NBDNJ and 19029 do not antagonize and appear to potentiate the anti-SARS-CoV-2 activity of Remdesivir. During the preparation of this manuscript, reports were made describing the effectiveness of NBDNJ in suppressing SARS-CoV-2 (ICGEB-FVG-5-Trieste) in Vero E6 cells, with an IC50 of 46 μM (Rajasekharan et al., 2020) and a discussion of the use of alpha glucosidase inhibitors in treating coronavirus infections (Williams & Goddard-Borger, 2020). Our results are consistent with these reported data. How our contribution adds to their findings is considered in the Discussion.

## METHODS

### Cells and Virus

Human lung carcinoma A549^ACE2^ cells were maintained in DMEM made 10% FBS (Weston et al., 2020). SARS-CoV-2 (USA-WA1/2020 strain) was deposited by the Centers for Disease Control and Prevention and obtained through BEI Resources (NR-52281). icSARS-CoV-2-mNG (Xie et al., 2020), which expresses a neon green from ORF7b, was provided by Dr. Pei-Yong Shi, University of Texas Medical Branch, Galveston, TX. Both viruses were propagated in Vero-E6 cells.

Human lung (carcinoma) A549^ACE2^ cells were seeded at a density of 10,000 cells per well. After one day, they were either left uninfected or infected with SARS-CoV-2 at an moi of 0.5. After incubation for 1 hour, media was removed and left in complete media made 1.0% DMSO (Fig.2A), or incubated with complete media and compound in 1% DMSO. After 48 hours, cells were stained with DAPI, and in the case of SARS-CoV-2 (USA-WA1/2020 strain) with anti nucleocapsid antibody. Images (20X) read in the blue filter to detect DAPI staining nuclei and the green filter to detect the viral nucleocapsid

**Figure 2:**
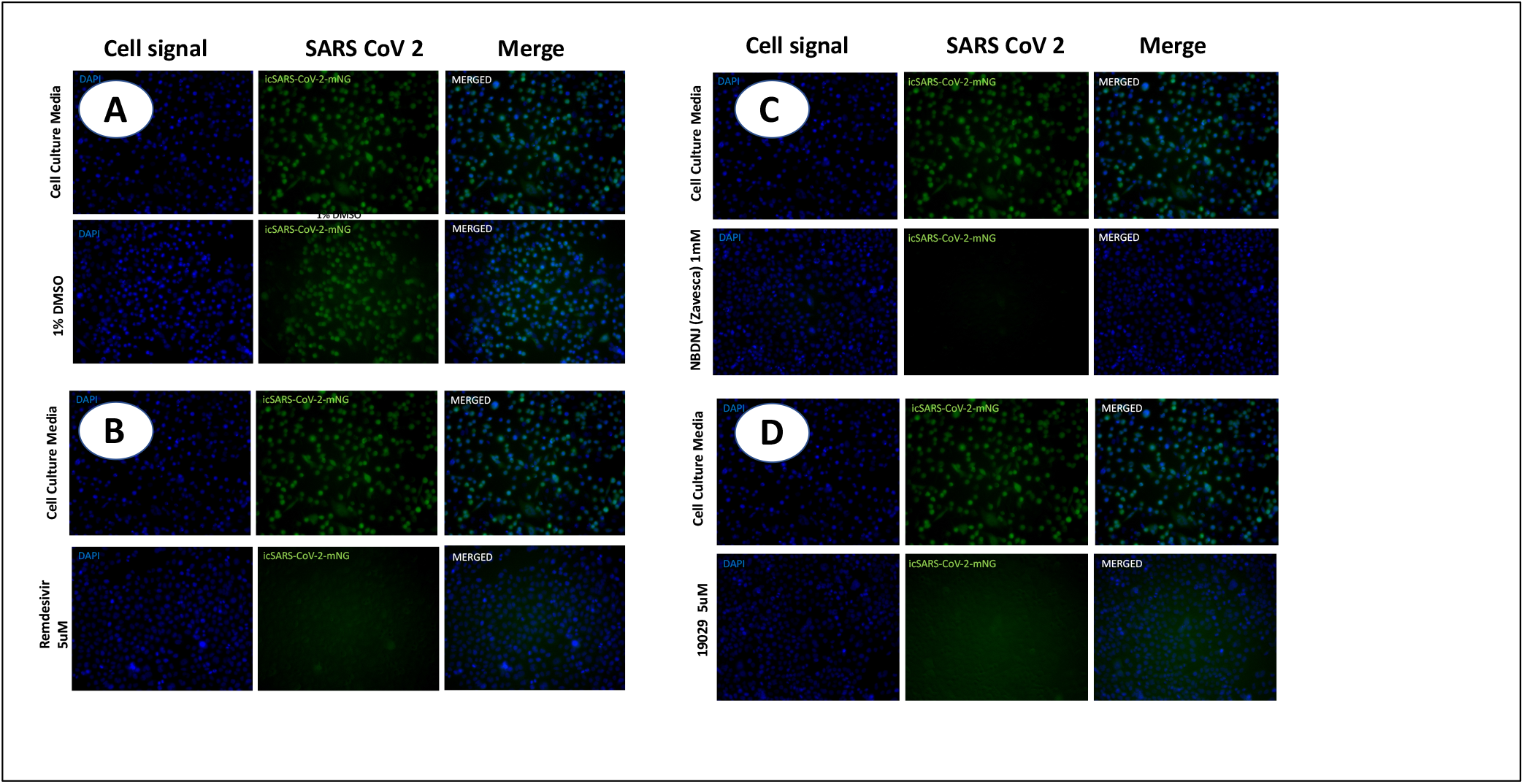
Infection with SARS-CoV-2 and repression with Remdesivir and iminosugars. Human lung (carcinoma) A549^ACE2^ cells, grown in DMEM and 10% FBS, were seeded at a density of 10,000 cells/well and, after one day, infected with infected with icSARS-CoV-2-mNG, at an moi of 0.5 in the absence and presence of varying concentrations of drugs. After incubation for 1 hour, media was removed and left in media made 1.0% DMSO (Fig.2A), or with the 5uM Remdesivir (Fig. 2B), 1 mM NBDNJ (Fig 2C) or 5uM 19029 (Fig. 2D) in 1.0% DMSO for 48 hours. Cells were stained with DAPI, and images (20X) read in the blue filter to detect DAPI staining nuclei and the green filter to detect the icSARS-CoV-2-mNG.

### Chemicals

Remdesivir was purchased from Ambeed, Inc. Iminosugars NBDNJ and BSBI 19029 to suppress SARS-CoV-2 replication in tissue culture, NBDNJ was obtained commercially, and 19029 was synthesized by starting from tert-butyl-dimethyl-silyl (TBDMS) protected bromohexyl-alcohol as in Du et al. (2013).

## RESULTS

To assess the ability of iminosugars NBDNJ and BSBI 19029 to suppress SARS-CoV-2 replication in tissue culture, NBDNJ was obtained commercially, and 19029 was synthesized by starting from tert-butyl-dimethyl-silyl (TBDMS) protected bromohexyl-alcohol as in Du et al. (2013). Their structures are shown in Fig. 1. It was of interest to test 19029, because although NBDNJ (also called Miglustat/Zavesca) was shown to be effective in managing Gauche Disease (Cox et al., 2000) and is currently approved for use for its management in people (Butters et al., 2005) the dose at which it is used clinically is lower than has been historically needed for antiviral suppression. (Higher dosing has been associated with diarrhea.) 19029 is a N-alkylated derivative of DNJ and is much more potent than NBDNJ and has been modified as a prodrug that would avoid inhibition of gastrointestinal glucosidases responsible for the GI distress (Du et al., 2013).

For antiviral analysis, A549^ACE2^ cells and both wild type SARS-CoV-2 and icSARS-CoV-2-mNG (Xie et al., 2020) were used. A549^ACE2^ cells were seeded at a density of 10,000 cells/well and incubated with icSARS-CoV-2-mNG at an moi of 0.5 in the absence and presence of varying concentrations of NBDNJ, 19029 or Remdesivir. 48 hours after infection, cells were fixed and the amount of SARS-CoV-2 was detected by imaging with a Perkin Elmer EnVision plate reader. SARS-ncCoV-2/USA/WA1/2020 was detected by staining with antibody to nucleocapsid. icSARS-CoV-2-mNG is a fully infectious virus expressing a neon green reporter expressed from ORF7b and thus replication can be easily monitored as shown in (Fig. 2A).

As expected, 5 μM Remdesivir almost completely suppressed all detectable SARS-2 signal (Fig. 2B). Encouragingly, NBDNJ and 19029 both also displayed significant antiviral activity, repressing most of the SARS-CoV-2 signal at 1000 μM and 5 μM, respectively (Fig. 2C & D, respectively). CC_50_s were not detected specifically measured in this assay, but in parallel experiments no toxicity was seen at >10 mM of NBDNJ (not detected) or >100 μM 19029.

The experiments were repeated with varying concentrations of each compound and as expected, shown in Fig. 3,

**Fig. 3.**
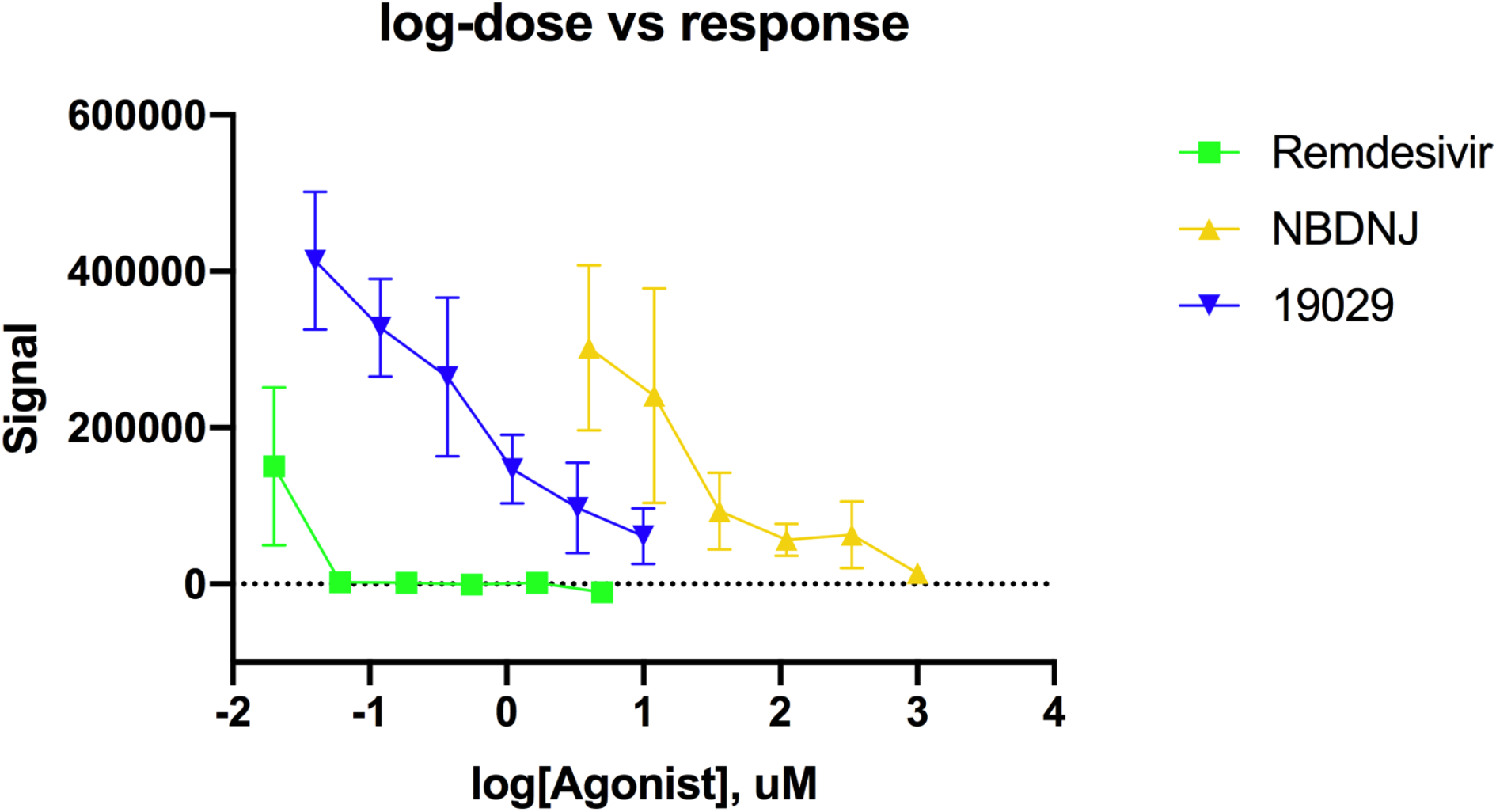
Dose response analysis of Remdesivir and the imino sugars in suppressing icSARS-CoV-2-mNG in tissue culture. Human lung (carcinoma) A549^ACE2^ cells were infected with icSARS-CoV-2-mNG in the absence and presence of vary concentrations of drugs, and after 48 hours, fixed and stained with DAPI, and images (20X) read in the blue filter to detect DAPI staining nuclei and the green filter to detect the icSARS-CoV-2-mNG. Fluorescence was quantified.

Remdesivir repressed icSARS-CoV-2-mNG in a dose responsive manner, with an IC_50_ estimated to be ~2.0 nM, in this assay. Both NBDNJ and 19029 also repressed icSARS-CoV-2-mNG in a dose responsive, non cytotoxic, with an IC_50_ of 17 μM and IC_90_ of 130 μM for NBDNJ and an IC_50_ of 450nM and IC_90_ of 4 μM for 19029.

Since Remdesivir is already in use for the management of SARS-CoV-2 infection (Al-Abdouh et al., 2021), we also wanted to determine if combinations of the 19029 or NBDNJ with Remdesivir were also effective. Fig. 4 shows graphically that the combination of 0.037 μM of NBDNJ or (Fig. 5) 0.04 μM 19029 with 0.002 μM Remdesivir almost completely eliminated the SARS-CoV-2 (USA-WA1/2020) signal. These data show there is no evidence of antagonism between these drugs. Ideally, we would like to have determined if NBDNJ or 19029 potentiated the effectiveness of Remdesivir. Although these results are encouraging, more work is needed to make conclusive determinations about specific dosing and combination effects.

**Fig. 4.**
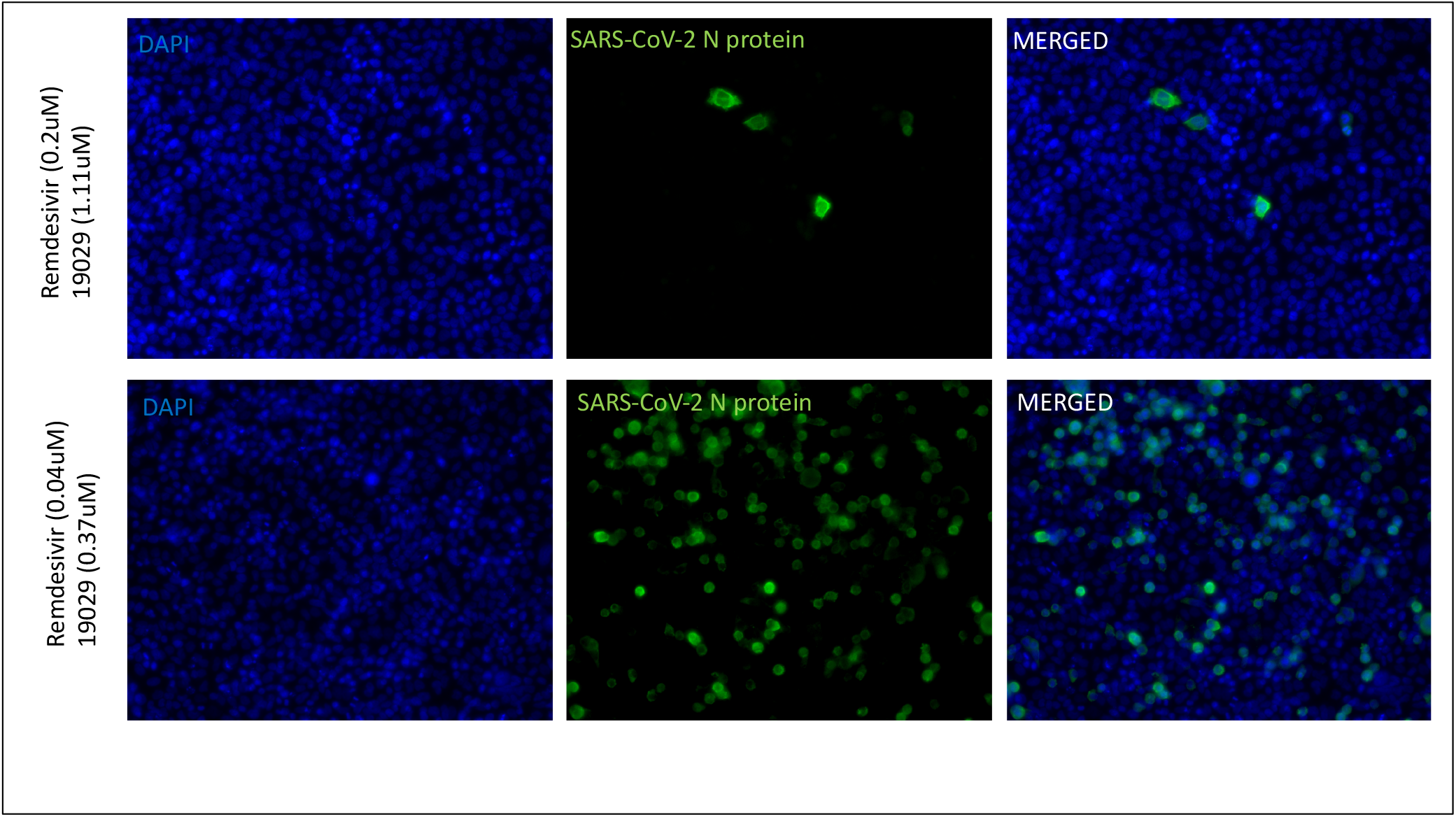
Combinations of NBDNJ and Remdesivir to suppress SARS-CoV-2. Human lung (carcinoma) A549^ACE2^ cells were infected with SARS-CoV-2 (USA-WA1/2020 strain) in the absence and presence of vary concentrations of drugs, and after 48 hours, stained with anti nucleocapsid antibody and images (20X) read in the blue filter to detect DAPI staining nuclei and the green filter to detect the viral nucleocapsid Fluorescence images are shown.

**Fig. 5.**
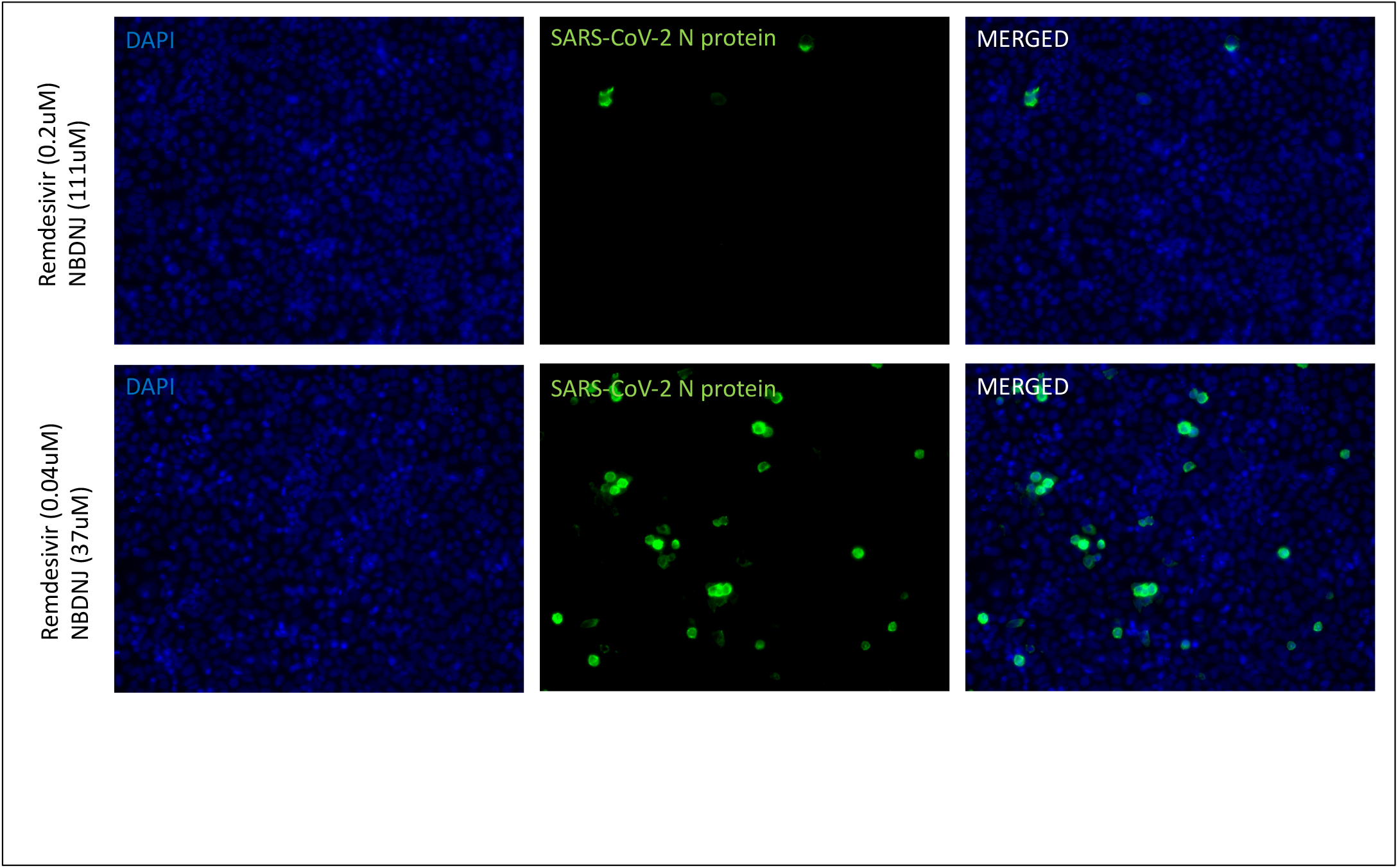
Combinations of 19029 and Remdesivir to suppress SARS-CoV-2. Human lung (carcinoma) A549^ACE2^ cells were infected with SARS-CoV-2 (USA-WA1/2020 strain) and incubated with the indicated combinations of drugs, as in Fig. 4. Staining and fluorescence is as in Fig. 4. Fluorescence images are shown.

## DISCUSSION

Although the precise concentrations of drugs used here may not directly correspond to concentrations needed in vivo, the results here are provocative. We provide tissue culture evidence that the iminosugars NBDNJ and 19029 are effective inhibitors of SARS-CoV-2. Both authentic and recombinant SARS-CoV-2 were used in our study. The use of NBDNJ to repress SARS-CoV-2 in tissue culture systems was recently reported by Walls et al. (2020). Thus, our work confirms those results, and, importantly, also shows the effectiveness of 19029, an imino sugar that is more potent and designed to cause fewer gastrointestinal events which resulted from inhibiton of intestinal glucosidases, than does high dose NBDNJ (Du et al., 2013). These imino sugars are ER glucosidase inhibitors. ER Glucosidases are necessary for ACE2 and SARS-1 Spike glycoprotein maturation. Since SARS-2 also uses ACE2 as its cell entry receptor, and its Spike polypeptide is heavily N-glycosylated, it is likely that this is also the basis of the drug’s SARS-2 antiviral activity.

Remdesivir was shown to have antiviral activity against SARS-CoV-2 (USA-WA1/2020) and icSARS-CoV-2-mNG, and this is not surprising, since it has established anti-SARS-CoV-2 (Wang et al., 2020; Simonis et al., 2021; Spinner et al., 2021). Given the imino sugar’s mechanism of action, it seemed likely that the imino sugars would complement, perhaps potentiate, the antiviral activity of Remdesivir. In our initial analysis, there is no antagonism observed between the iminosugars and Remdesivir and there may even be potentiation, although more work to establish this is needed.

There is an urgency in evaluating promising existing antiviral strategies to manage SARS-CoV-2. Since NBDNJ is already a drug approved for human use, it is possible that evaluation of its value in managing SARS-CoV-2 could be explored in people fairly rapidly. That it does not antagonize and may even potentiate Remdesivir, which has been shown to be effective, but of limited therapeutic value, is particularly exciting. Remdesivir is given parenterally, however. I.V. formulations of NBDNJ are certainly possible, and other parenteral use of NBDNJ (experimentally in animals) has been done. We also have preliminary evidence that NBDNJ and 19029 may complement Favipiravor. This would also be important, since Favipiravor has also been reported to have activity against SARS-2 and is orally available, and in human use (Driouich et al., 2021).

19029 is a more potent derivative of NBDNJ and in its prodrug form, can pass the GI track with a minimum of GI glucosidase inhibition and GI distress. Although GI upset associated with long term high dose NBDNJ might not be a problem for the short-term use anticipated in management of COVID-19, it is reassuring to have a second-generation compound such as 19029. Needless to say, more work is needed. Preclinically, animal studies are indicated, and clinically, careful consideration of the usefulness of NBDNJ alone and in combination with Remdesivir may be justified, and this should include proper dose finding.

